# Compensation of the effects of temperature on a motor system in the crab, *Cancer borealis*

**DOI:** 10.64898/2026.06.23.734078

**Authors:** Kathleen Jacquerie, James M. DiMartino, Ananya Dalal, Ji Zeng, Eve Marder

## Abstract

Rising ocean temperatures challenge ectothermic animals to maintain essential behaviors such as movement and feeding. We asked how a complete neuromuscular pathway preserves function when every component process responds differently to warming. In the pyloric system of the crab *Cancer borealis*, we simultaneously recorded motor nerve activity, muscle membrane potential, and contraction. Warming preserved rhythmic nerve activity and excitatory junctional potentials, but contraction declined and failed first. Fixed low-frequency stimulation, mimicking cold-temperature motor output, resulted in reduced contraction at warm temperatures, whereas higher-frequency stimulation, mimicking warm-temperature motor output, partially restored contraction. Warming hyperpolarized muscle fibers, moving them farther from contraction threshold, but also reduced input resistance, which together limited over-excitability. However, high-potassium stimulation revealed that the muscle contractile machinery remained functional. Thus, warming acts differently across levels, and overlapping compensatory mechanisms help preserve neuromuscular function across a wide range of temperatures.

**Significance statement:** Cold-blooded animals that live in climates with significant seasonal changes in ambient temperature must have myriad mechanisms to function over a wide range of environmental conditions. We explore the effects of temperature at multiple levels of organization within the stomatogastric system of the crab, *Cancer borealis*. We find a series of compensatory mechanisms that cooperatively help maintain stable function despite the fact that the motor patterns, neuromuscular junctions and muscle functions are all differently temperature dependent.

## Introduction

Crustaceans, such as lobsters and crabs, are poikilotherms and some species inhabit environments with large daily and seasonal temperature variations. In their New England Atlantic Ocean habitat, the crab *Cancer borealis* experiences temperatures ranging from ∼3°C to ∼25°C (Marder *et al*., 2015; ‘NOAA’, 2026). Despite temperature’s profound influence on neuronal and muscular components, these animals maintain life-critical processes across this range. Understanding how temperature compensation cooperates across different organizational levels presents an intriguing challenge. Specifically, we ask whether temperature changes affect motor pattern production, the synapse between the motor neuron and the muscles they innervate, and nerve-evoked muscle movement coordinately, or does temperature affect these processes to a greater or a lesser degree?

Temperature affects both passive and active properties of neuromuscular junctions across a wide range of species, both vertebrates and invertebrates. Much work has been done in species like the frog (*Rana pipiens*) (Katz and Miledi, 1965), crayfish (*Astacus leptodactylus, Procambarus clarkii*) (Harri and Florey, 1977; Fischer and Florey, 1981; White, 1983), crabs (*Carcinus maenas*, *Cancer pagurus*, *Ocypode ceratophthalma*) (Florey and Hoyle, 1976; Stephens and Atwood, 1983; Philip J. Stephens, 1985; Hyde *et al*., 2015), barnacles (Dipolo and Latorre, 1972), rat diaphragm (Ward, Crowley and Johns, 1972; Head, 1983), drosophila (Ormerod, Scibelli and Littleton, 2022). In general, warming shortens synaptic delay and often reduces the amplitude and temporal summation of the excitatory junctional potentials (EJPs). Warming can also increase miniature end-plate potential frequency (Ward, Crowley and Johns, 1972). Warming alters passive membrane properties. It commonly hyperpolarizes the resting membrane potential, decreases input resistance, reducing synaptic integration (Fischer and Florey, 1981; Jacquerie *et al*., 2026). At the mechanical level, warming weakens nerve-evoked contractions (Foldes *et al*., 1978; Fischer and Florey, 1981; Thuma *et al*., 2013). The excitation-contraction threshold in the crayfish opener muscle is relatively temperature independent (Fischer and Florey, 1981). Together, these studies indicate that warming can challenge neuromuscular systems through combined effects on synaptic transmission, membrane properties, and force production.

Many studies have also examined the effects of temperature on motor pattern generation. The stomatogastric nervous system of crabs and lobsters, particularly the pyloric circuit involved in food filtering has been extensively studied. In this system, warming generally increases pyloric rhythm frequency while preserving the relative phase relationships of the motor pattern over a broad permissive temperature range. At higher temperatures, however, the rhythm becomes irregular or fails, a phenomenon often referred to as a “crash” (Tang *et al*., 2010, 2012; Marder *et al*., 2015). In *Cancer borealis*, pyloric rhythm frequency approximately doubles between 11°C and 21°C (Tang *et al*., 2010, 2012; Pereira *et al*., 2014; Soofi *et al*., 2014; Marder *et al*., 2015; Städele, Heigele and Stein, 2015; Haddad and Marder, 2018; Kushinsky, Morozova and Marder, 2019; Powell *et al*., 2021; Städele and Stein, 2022; Hampton, Kedia and Marder, 2024; Schapiro *et al*., 2024). These observations raise a central question: how does the neuromuscular system respond when motor pattern frequency increases with temperature?

Similar principles have been described in other motor pattern generators; in the ventilatory central pattern generator of the locust *Locusta migratoria* (Armstrong *et al*., 2006), in the swimming pattern of the *Xenopus laevis* tadpoles (Sillar and Robertson, 2009), and in the locomotor output of the larval *Drosophila* (Barclay, Atwood and Robertson, 2002). Warming initially speeds motor rhythms, but excessive heat challenges circuit stability and can ultimately lead to failure.

Motor pattern generation and neuromuscular transmission have usually been studied separately. Here, we instead examine multiple components of the neuromuscular system together to ask whether compensatory or cooperative mechanisms emerge across levels of organization. The closest related studies are those on the crustacean heart (Worden *et al*., 2006; Kushinsky, Morozova and Marder, 2019), where warming increases beat frequency while contraction amplitude eventually declines, and the work of Thuma and colleagues on the pyloric muscle of the spiny lobster *Panulirus interruptus*, showing that contraction fails before the pyloric rhythm itself as temperature rises (Thuma *et al*., 2013). Here, we study both the pyloric motor pattern and its target muscles. This work reveals how cooperative changes across the motor circuit, neuromuscular synapse, and muscle help sustain motor output during temperature challenge.

## Results

### Pylor contraction failed before motor pattern at high temperatures

Figure 1A shows the stomatogastric nervous system (STNS) and several pyloric muscles in *Cancer borealis* (Weimann, Meyrand and Marder, 1991; Jacquerie *et al*., 2026). The stomatogastric ganglion (STG) contains the motor neurons that generate the pyloric rhythm: one lateral pyloric neuron (LP), several pyloric neurons (PY), and two pyloric dilator neurons (PD). Together, these neurons produce the triphasic pyloric rhythm that drives movements involved in filtering food in the foregut. The LP and PY neurons constrict the pylorus, while PD neurons dilate the pylorus (Marder and Bucher, 2007). The axons of PD, LP and PY travel through the lateral ventricular nerve (lvn) before branching to innervate specific muscles. Here, we illustrate three LP-innervated muscles, cpv4, cpv6, and p1, and one PY-innervated muscle, p2, all of which are glutamatergic. Figure 1B shows the experimental configuration used for multilevel recordings: the lvn was recorded extracellularly with a suction electrode, muscle fibers were impaled with intracellular electrodes, and the muscle output produced by the three LP-innervated muscles was measured with a force-displacement transducer.

**Figure 1.**
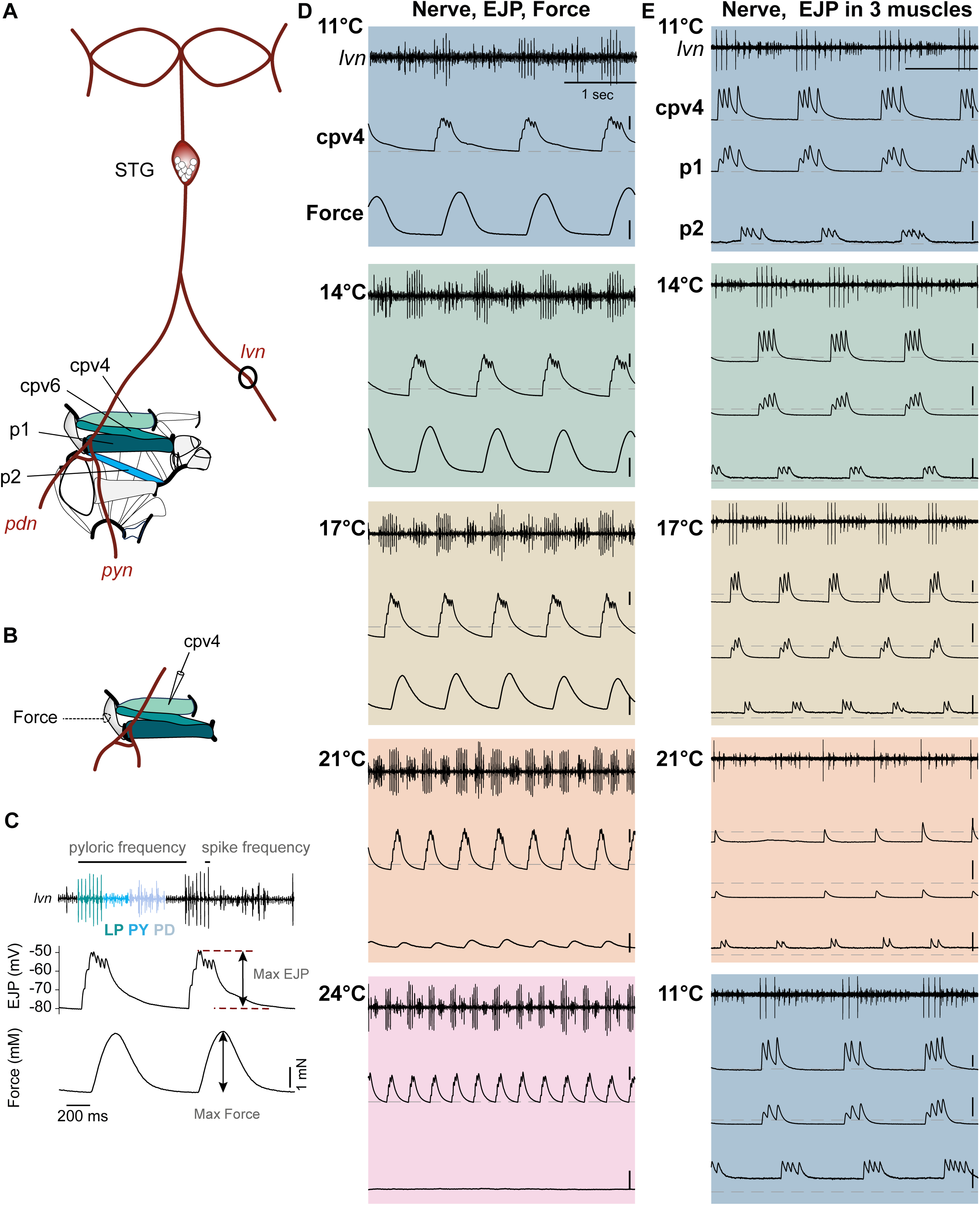
Temperature-sensitivity of the different levels of organization at the neuromuscular system in the crab *Cancer borealis.* (A) Schematic of the stomatogastric system, showing the nerve diagram, the stomatogastric ganglion (STG), innervating the pyloric (p) and cardiopyloric (cpv) muscles via the lateral ventricular nerve (lvn), pyloric dilator nerve (pdn), and pyloric nerve (pyn), according to the nomenclature of (Maynard and Dando, 1974; Weimann, Meyrand and Marder, 1991). Only the left side of the stomach is shown, with the posterior oriented downward. (B) Schematic of the in vitro preparation where muscles are dissected and kept connected to their motor nerves. The force-displacement transducer was attached to the cartilage linked to the three LP-innervated muscles: cpv4, cpv6, and p1. (C) Example of simultaneous multi-level recordings during rhythmic pyloric activity, showing lvn activity, STG-evoked excitatory junction potentials (EJPs) recorded intracellularly in cpv4, and the corresponding force produced by the three LP-innervated muscles (cpv4, cpv6, and p1). Maximal EJP amplitude and maximal force are indicated. (D) Simultaneous recordings of lvn activity, cpv4 EJP, and muscle force during warming. While rhythmic nerve activity and EJPs were maintained from 11 to 24°C, force progressively declined and failed first. (E) Simultaneous recordings of lvn activity and intracellular responses from two LP-innervated muscles (cpv4 and p1) and one PY-innervated muscle (p2) during warming. Dashed lines indicate the resting membrane potential at 11°C (D: cpv4, V_m_ = −75 mV; E: cpv4, V_m_ = −67 mV; p1, V_m_ = −69 mV; p2, V_m_ = −101 mV). Vertical scale bars for intracellular recordings are 10 mV; force scale bar, 1 mN.

Previous work has shown that in *Cancer borealis*, *Carcinus maenas*, and *Homarus americanus*, motor neuron burst frequency increases with temperature while phase relationships are largely maintained, until the rhythm crashes at high temperatures (Tang *et al*., 2010; Soofi *et al*., 2014; Stein *et al*., 2023; Schapiro *et al*., 2024; Carrier *et al*., 2026). Here, we extend these studies by combining recordings of motor nerve activity, muscle fiber membrane voltage, and muscle contraction driven by the spontaneous in vitro pyloric rhythm. Figure 1C shows an example of this multilevel recording. The triphasic pyloric rhythm can be seen on the lvn as the successive activity of LP, PY, and PD. In cpv4, each motor neuron action potential evokes an EJP, and the muscle integrates these depolarizations to generate contraction.

Figure 1D illustrates the effect of increasing saline temperature from 11°C to 24°C. As temperature increases, the pyloric rhythm increases frequency, as shown by the shorter cycle period between LP bursts. In cpv4, EJP amplitude remains relatively large and continues to follow the phase-compensated motor neuron input. However, although contraction remains strong from 11°C to 17°C, it weakens at 21°C and disappears at 24°C. These data show that contraction fails at lower temperatures than motor neuron bursting or the muscle depolarization itself. This observation is consistent with earlier work in the spiny lobster *Panulirus interruptus*, in which pyloric network activity remained rhythmic between 9°C and 15°C, while the p1 muscle stopped at 15°C (Thuma *et al*., 2013).

Figure 1E shows simultaneous recordings of nerve activity and EJPs in cpv4, p1, and p2 in another animal. Each muscle depolarizes in response to motor neuron action potentials. At 21°C, however, the rhythm starts to crash, leaving only sparse action potentials that evoke small EJPs. After cooling, the rhythm recovers and EJPs are again maintained.

To quantify these temperature-dependent changes in pooled data from multiple preparations, we plotted the pyloric network activity and muscle responses as a function of temperature (Figure 2). The mean pyloric network frequency increased from 0.97 ± 0.06 Hz at 11°C to 2.26 ± 0.19 Hz at 23°C, with a significant overall effect of temperature (Linear Mixed Effect, LME, p = 1.44 × 10^−18^) and significant increases relative to 11°C at all warmer temperatures (Holm-corrected comparisons; Figure 2A). A Q_10_ value is used to characterize the temperature sensitivity of the frequency, as previously described (Tang *et al*., 2010). The mean Q_10_ of the pyloric frequency is 1.96 ± 0.11 (n = 14). This indicates that, over this temperature range, pyloric frequency approximately doubles for a 10°C increase. The mean LP duty cycle was largely maintained from 11°C to 17°C, but decreased at warmer temperatures, from 20.7 ± 1.8% at 11°C to 13.4 ± 3.1% at 23°C; this corresponded to a significant overall effect of temperature (LME, p = 4.5 × 10^−6^), with Holm-corrected comparisons showing significant decreases at 20°C and 23°C relative to 11°C (Figure 2C). These data are consistent with previous experiments (Tang *et al*., 2010, 2012; Schapiro *et al*., 2024).

**Figure 2.**
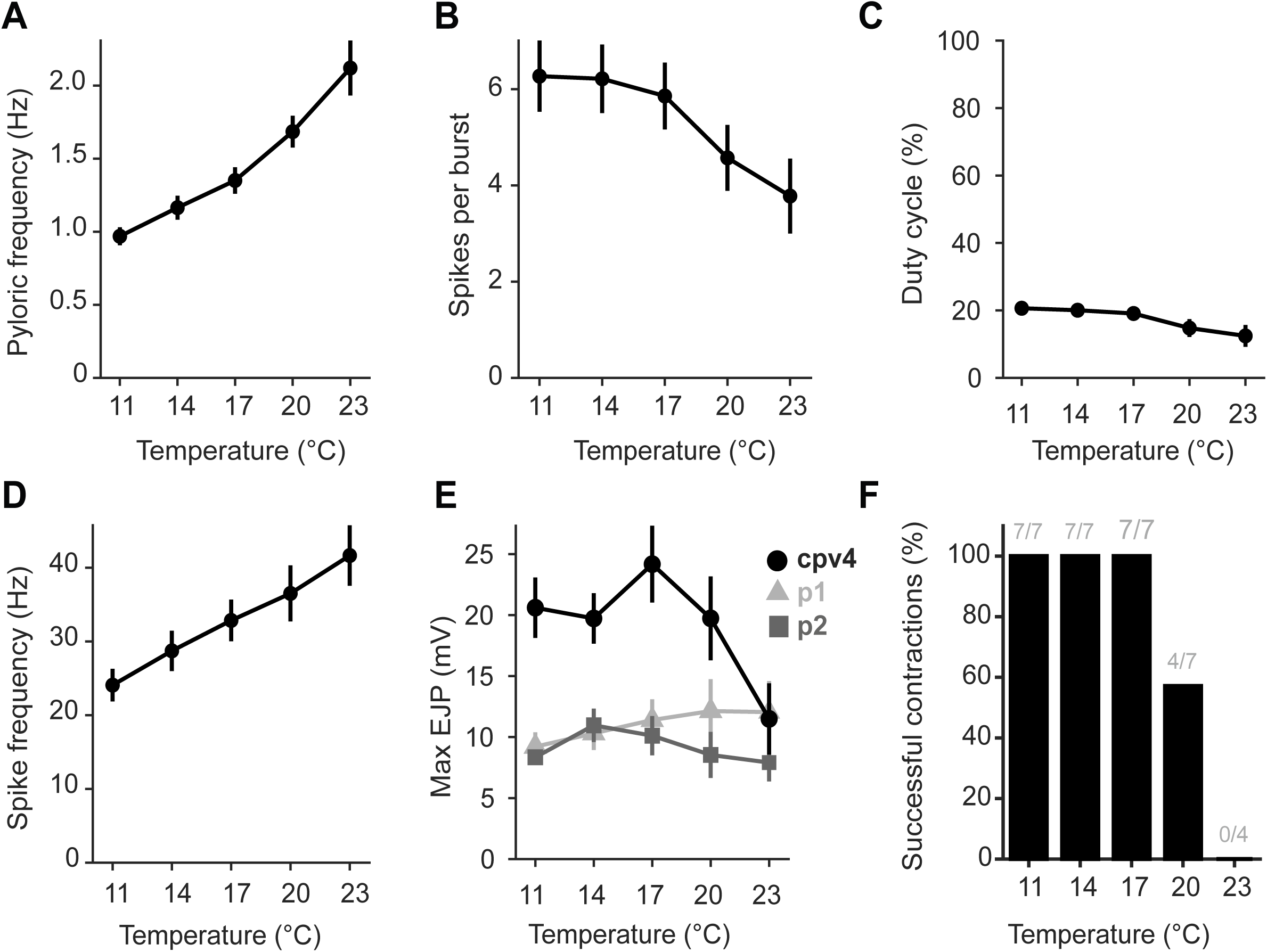
Temperature sensitivity of pyloric rhythm, STG-evoked EJPs, and muscle contraction. Contraction fails before pyloric rhythmic activity at high temperature. (A-D) Temperature dependence of pyloric rhythm parameters (illustrated in Figure 1C) averaged across preparations on LP: (A) pyloric frequency, (B) spikes per burst, (C) duty cycle (phase of LP activity divided by pyloric cycle period), and (D) spike frequency within the LP burst. (E) Maximum EJP amplitude as a function of temperature in three muscles: cpv4 (black circles), p1 (light gray triangles), and p2 (dark gray squares), across preparation. (F) Percentage of successful contractions as a function of temperature. A successful contraction was defined as a preparation in which pyloric nerve activity remained rhythmic and contraction amplitude was at least 15% of its value at 11°C.

We are interested in the coordination between the motor neuron input and the output of the neuromuscular system, as muscle depolarization and contraction are driven by both the number of spikes and the spike frequency within each burst. The mean number of spikes per burst remained close to 6 during the first part of the temperature ramp, from 11°C to 17°C, and then decreased at warmer temperatures, reaching 3.9 ± 0.75 spikes per burst at 23°C; this decrease was supported by a significant overall effect of temperature (LME, p = 6.5 × 10^−8^), with significant reductions at 20°C and 23°C relative to 11°C in Holm-corrected comparisons (Figure 2B).

The variability between animals is large and is consistent with the data reported by Tang et al., 2012. In contrast, the mean LP spike frequency increased monotonically from 24.1 ± 2.2 Hz at 11°C to 44.1 ± 3.9 Hz at 23°C, with a significant overall effect of temperature (LME, p = 2.27 × 10^−10^) and significant increases relative to 11°C at all warmer temperatures (Holm-corrected comparisons; Figure 2D). The mean Q_10_ of the spike frequency is 1.6 ± 0.1 (n=14). Interestingly, the average max EJP amplitude was not uniformly affected by temperature across muscles. While p1 and p2 showed relatively small changes over the tested temperature range, cpv4 displayed a marked non-monotonic profile, with amplitudes maintained or slightly increased at intermediate temperatures before decreasing at warmer temperatures. Consistent with this muscle-specific profile, a linear mixed-effects model showed a significant effect of muscle identity (p = 4.3 × 10^−6^) and a significant muscle and temperature interaction (p = 0.012), but no main effect of temperature across all muscles (p = 0.88). This high-temperature decline in cpv4 EJP amplitude occurred over the same temperature range in which LP spike number decreased, suggesting that changes in motor burst structure may contribute to the reduced compound depolarization.

Figure 2F summarizes the ability of the stomatogastric system to generate a pyloric motor pattern that effectively drives contractions across temperatures. Across 7 preparations, contractions were maintained at lower temperatures, but at 20°C, 3 preparations still exhibited rhythmic motor input without producing contraction. In the 4 remaining preparations that still contracted at 20°C, a further increase of 3°C abolished muscle contraction, although rhythmic nerve activity persisted.

Together, these results show that warming does not first abolish rhythmic motor activity or muscle depolarization but instead disrupts the conversion of rhythmic motor input into effective mechanical output. This suggests that temperature-compensation mechanisms operating at individual levels of the neuromuscular system are insufficient to preserve integrated mechanical function at elevated temperatures.

### Increasing stimulation frequency partially rescues contraction at high temperatures in an isolated neuromuscular preparation

To isolate the contribution of motor input parameters from the temperature-dependent changes in network activity, we severed the lvn and stimulated it with controlled burst trains while varying saline temperature independently. Each train consisted of 20 bursts of 8 spikes, a spike number chosen to match the mean LP burst size observed in our recordings (Figure 1C) and consistent with values reported by Tang et al. (2012). We designed four stimulus patterns, “LP 6”, “LP 11”, “LP 16”, and “LP 21”, to mimic the LP neuron activity recorded at 6, 11, 16, and 21°C, respectively, varying both pyloric frequency (0.5 to 2 Hz) and intraburst spike frequency (20 to 60 Hz) (Figure 3A, top).

**Figure 3.**
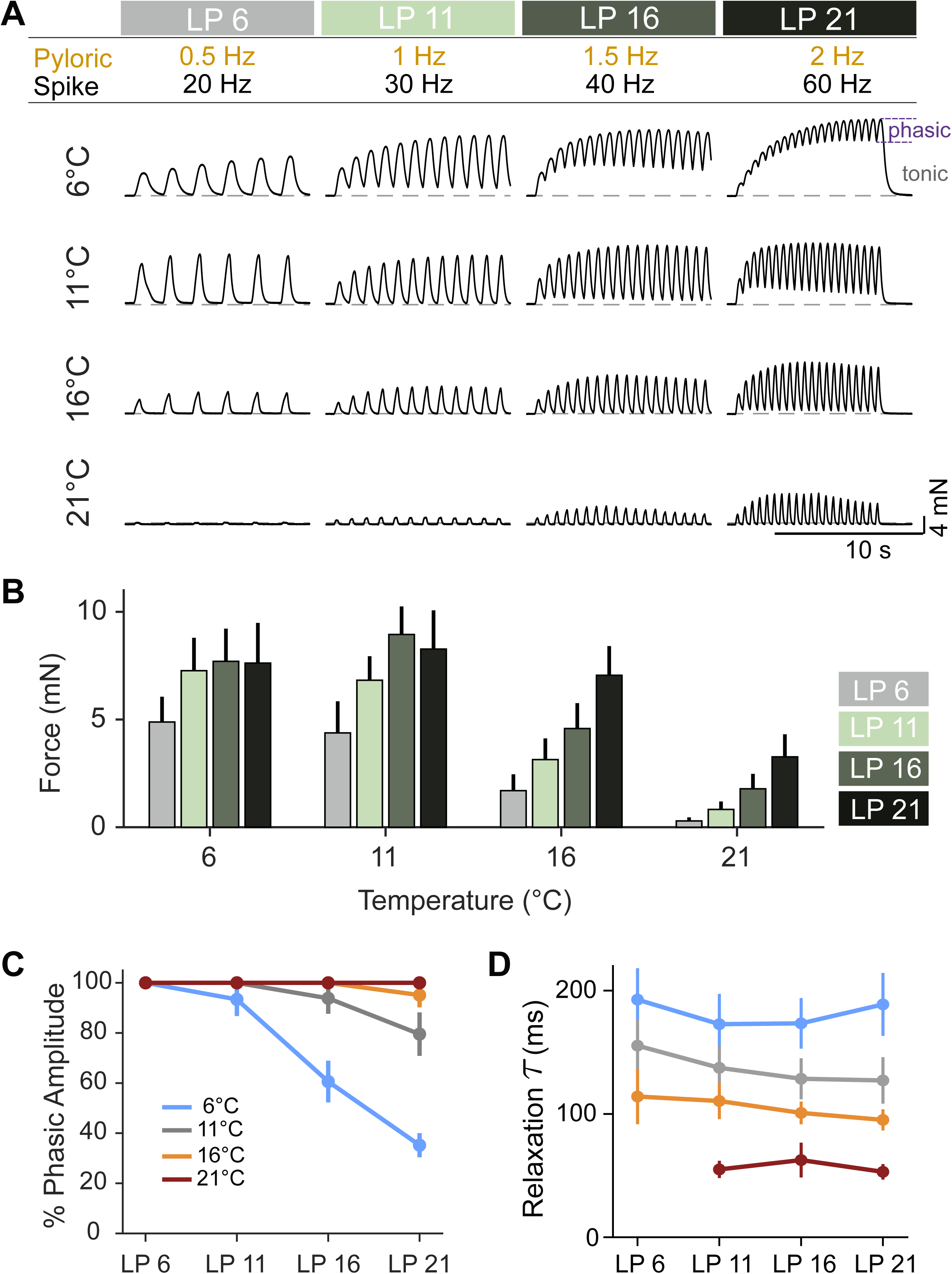
Nerve-evoked contraction decreases with temperature and is partially rescued by increasing stimulation frequency. (A) Contractions are evoked in LP-innervated muscles cpv4, cpv6, and p1, by stimulating the lvn in an isolated preparation. Stimulation trains reproduced stereotyped LP motor neuron firing patterns at different temperatures (LP 6, LP 11, LP 16, and LP 21). Pyloric frequency denotes the inverse of the period between two bursts, and spike frequency denotes within-burst firing frequency. Spike number is fixed at 8 per burst, within 20 bursts per train. Representative raw force traces are shown for each temperature. Dashed line, baseline tension. (B) Maximal force evoked by each stimulation pattern as a function of temperature (n=7). Bars show mean ± SEM across preparations. Higher stimulation frequencies (in darker green) increased force and partially rescued contraction at elevated temperatures. (C) Percent phasic is the phasic amplitude divided by the total amplitude of the contraction. (D) The time constant of relaxation decreased as temperature increased, while frequency of stimuli had little effect.

At a fixed stimulation pattern, muscle force depended strongly on temperature, with contraction amplitude decreasing at elevated temperatures (Figure 3A, by columns; LME main effect of temperature, p = 9.82 × 10^−5^, n = 7 animals). This is consistent with the temperature-dependent decline in contraction force reported by Thuma et al. (2013) for the p1 muscle of *Panulirus interruptus* under controlled nerve stimulation.

At a fixed temperature, stimulation pattern also affected the maximum force reached during the 20-burst train (Figure 3A, by rows; LME main effect of stimulation pattern, p = 0.039). The fastest pattern (LP 21) generally produced greater force than the slowest (LP 6) at every temperature tested.

At low temperatures (6°C, 11°C), high-frequency stimulation caused temporal summation of individual contractions: the tonic amplitude, defined as the minimum contraction to which the muscle relaxes between neuron bursts, increased, while the phasic amplitude, defined as the oscillation amplitude riding on this tonic baseline, decreased. This was clearly visible for LP 21 at 6°C (raw trace, Figure 3A) and it is consistent with the frequency-dependent contraction components described by Morris and Hooper (1997, 1998). This reduction of the phasic amplitude at cold temperature for a high frequency pattern is shown in Figure 3C for 7 animals, and demonstrates that the muscle cannot relax fast enough to follow high frequency discharge. Figure 3D supports that relaxation time constant considerably decreases with temperature, from 182 ms at 6°C to 57 ms at 21°C.

At 21°C, contraction was better preserved when the nerve was stimulated with the fastest LP-like pattern. LP21 produced larger contractions than both LP6 and LP11 at 21°C (Holm-corrected p = 0.019 for both comparisons), indicating that higher-frequency motor input partially compensated for the temperature-dependent loss of force. However, this rescue was incomplete: LP21 contractions at 21°C remained smaller than LP21 contractions at 11°C (Holm-corrected p = 0.019), and they also remained below the force produced by the physiologically matched LP11 pattern at 11°C (one-sided comparison, Holm-corrected p = 0.006).

These results suggest that the temperature-dependent increase in LP burst frequency observed in semi-intact preparation (Figure 1) provides partial functional compensation. At elevated temperature, a high-frequency LP-like pattern preserved more force than lower-frequency patterns, indicating that circuit-level increases in motor burst frequency can mitigate, but not fully prevent, the temperature-dependent decline in muscle output.

### Warming reduces the effectiveness of EJPs at the neuromuscular junction

During STG-evoked motor activity, contractions failed at high temperature while EJPs were still present, suggesting that the loss of mechanical output was not simply caused by the disappearance of synaptic input. To test how temperature affects neuromuscular transmission under controlled conditions, we cut the nerve and stimulated the lvn with paired pulses from 1, 5, 10, 20, 30, and 40 Hz across temperatures (11°C, 16°C, 21°C, and 26°C).

Paired-pulse stimulation increased EJP amplitude at all temperatures and frequencies (Figure 4A-B). EJP_2_ was significantly smaller at 21°C than at 11°C across all stimulation frequencies (Holm-corrected p ≤ 0.024 for all frequencies), showing that warming reduced the enhancement of the synaptic response produced by the second stimulus. At 26°C, spontaneous EJPs were observed, indicating spontaneous nerve discharge at this temperature.

**Figure 4.**
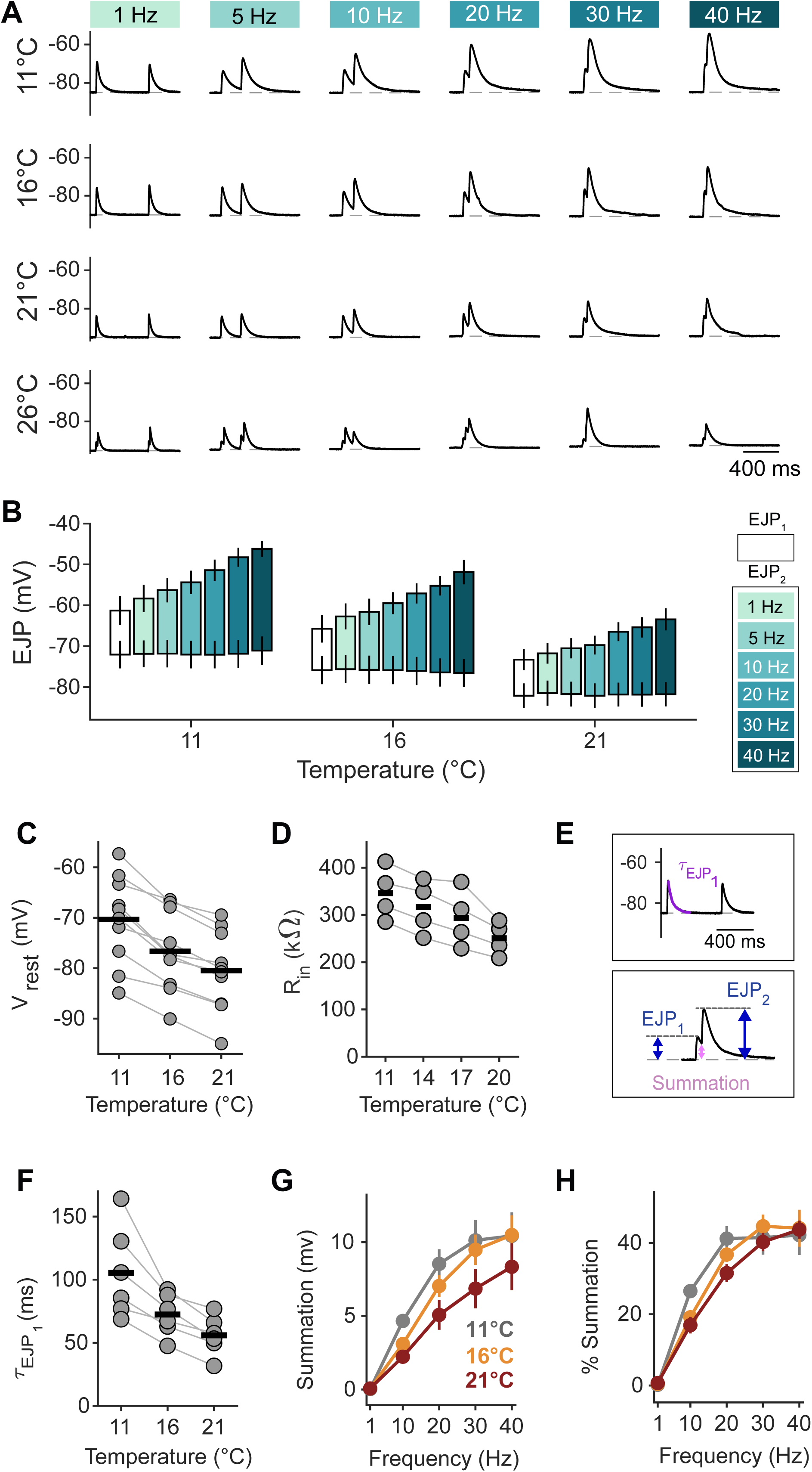
Warming reduces EJP amplitude and summation, hyperpolarizes cpv4, and decreases input resistance. (A) EJPs evoked in cpv4 by lvn stimulation with a paired-pulse protocol at 1, 5, 10, 20, 30, and 40 Hz. Representative traces are shown for each temperature. Dashed line is the resting membrane voltage (V_rest_). STG-evoked EJPs are observed at 26°C. (B) Floating-bar representation of EJP_1_ and EJP_2_ responses as a function of temperature and paired-pulse frequency. The lower bound represents the resting membrane voltage (V_rest_), and the upper bound represents the peak membrane voltage reached during the EJP. EJP_1_ is shown in white, and EJP_2_ is shown for each stimulation frequency. Bars show mean ± SEM for both V_rest_ and EJP peak voltage (7 fibers from 5 animals). (C) V_rest_ of cpv4 as a function of temperature (10 fibers from 6 animals). (D) Input resistance (R_in_) of cpv4 decreases as a function of temperature. (E) Feature of the paired-pulse experiments (F) Decay time constant of the EJP_1_ from the paired-pulse at 1 Hz. (G-H) Absolute summation and percent summation as a function of temperature and paired-pulse frequency. Percent summation was calculated as summation divided by EJP_2_ amplitude.

A linear mixed-effects model confirmed a strong main effect of stimulation frequency on EJP_2_ amplitude (p = 7.6 × 10^−15^), a significant main effect of temperature (p = 0.015), and no significant frequency × temperature interaction (p = 0.75). The absence of a significant interaction indicates that the frequency-dependent facilitation of EJP_2_ is preserved in shape across temperatures - warming shifts the amplitude curve downward uniformly rather than selectively abolishing high-frequency enhancement. Even at 21°C, high-frequency stimulation still produced larger EJPs than low-frequency stimulation (40 Hz vs 1 Hz at 21°C: Holm-corrected p = 5.5 × 10^−8^). However, this frequency-dependent compensation was incomplete: the second EJP at 40 Hz and 21°C was not significantly different from the second EJP at 10 Hz or 20 Hz at 11°C (Holm-corrected p = 0.64 and p = 0.18, respectively).

Thus, the temperature-dependent increase in motor burst frequency cannot fully rescue EJP amplitude at elevated temperatures. Several passive membrane changes likely contribute to this reduced synaptic efficacy. Resting membrane potential hyperpolarized progressively with warming (Friedman test, p = 0.0025, N = 10 fibers, n = 6 animals, Figure 4C), moving the membrane farther from the muscle excitation threshold. Input resistance calculated at steady-state also decreased with temperature (Friedman test, p = 0.0074, n = 4 animals, Figure 4D), meaning the same synaptic current produces a smaller voltage deflection. This is consistent with previous results on other stomach muscles (Jacquerie et al, 2026).

Finally, we measured the decay time constant of the first EJP and quantified summation as the residual depolarization from the first EJP remaining before the second EJP (Figure 4E). The EJP decay time constant shortened with warming (Friedman test, p = 0.0183, N = 6 fibers, n = 4 animals, Figure 4F), reducing the temporal overlap between successive EJPs. Because summation depends on how much residual depolarization remains when the next stimulus arrives, a shorter decay time constant is expected to reduce absolute summation. Consistent with this, summation increased with stimulation frequency, as expected from the shorter interval between paired pulses, but was reduced at warmer temperatures (Figure 4F). This agrees with previous work done in crayfish muscle (Fischer and Florey, 1981). When summation was normalized to EJP_2_ amplitude, percent summation changed little with temperature (Figure 4H), suggesting that warming mostly scaled down the EJP waveform rather than selectively reducing the relative contribution of summation.

### Contractile machinery remains functional at high temperatures

The loss of nerve-evoked contractions at high temperatures is partly explained by a decrease in the efficacy of neuromuscular transmission. However, the temperature sensitivity of the contractile machinery of the muscle itself is unknown and may also contribute to the loss of nerve-evoked contractions. To address this, we depolarized the muscles by applying saline solution with 10x the standard concentration of potassium for 40 s to elicit contractures at 11, 16, 21, and 26°C (Figure 5A).

**Figure 5.**
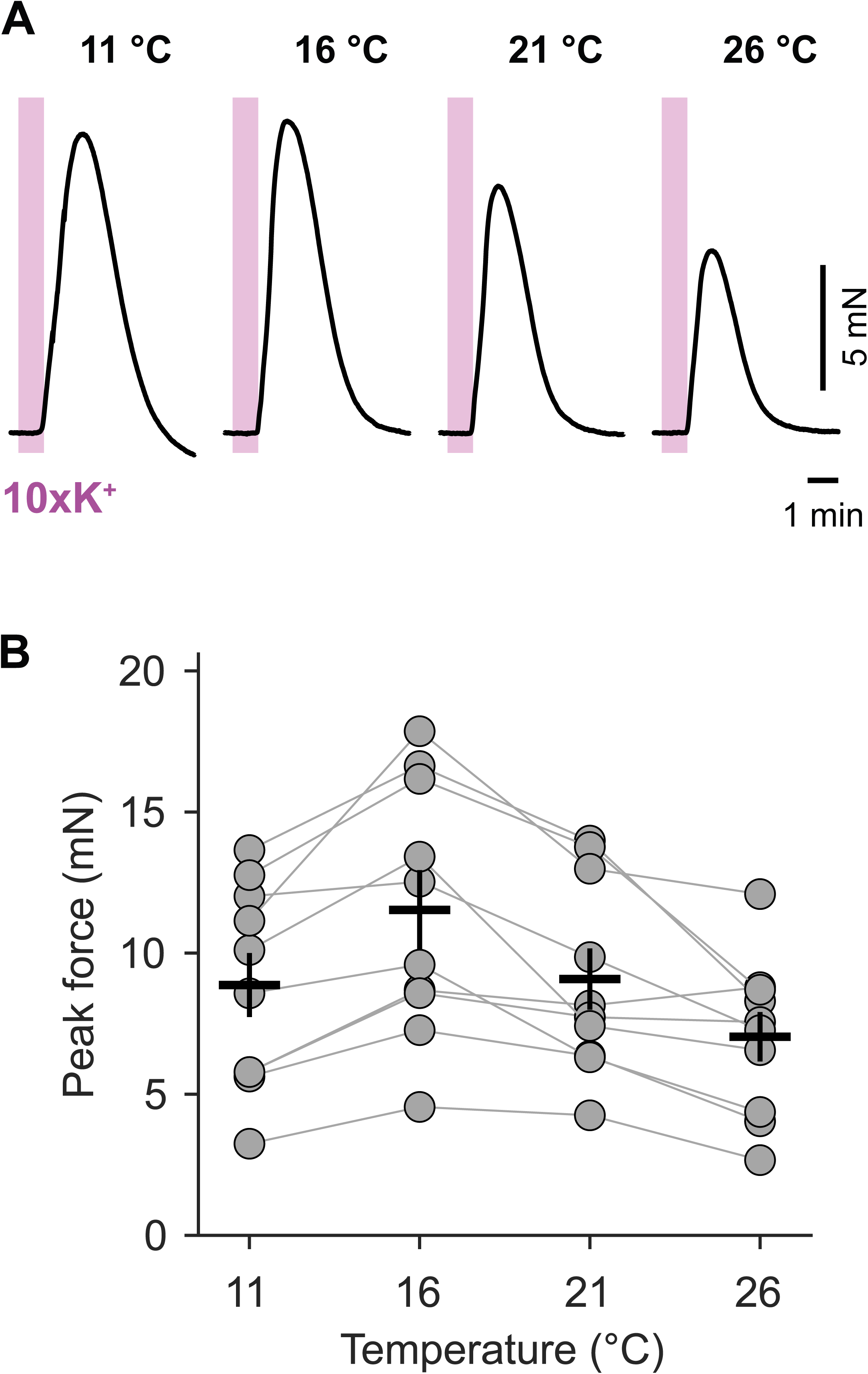
Contractile capacity of LP-innervated muscles is maintained across temperature. (A) Potassium-induced contractures in the LP-innervated muscles cpv4, cpv6, and p1. Contractures are elicited by applying saline containing 10 times the normal potassium concentration (10xK^+^) for 40 s with a flow rate of 25ml/min. Muscles were allowed to recover for 30 min between applications then temperature is increased at each application. (B) Maximal force generated by 10xK^+^ application as a function of temperature, shown as mean ± SEM across preparations (10 animals).

We measured the peak force produced during these contractures (n = 10 animals, Figure 5B). Temperature had a significant effect on contracture amplitude (Friedman test, p = 0.00023). Contracture amplitude increased moderately at 16°C compared to 11°C (Holm-corrected 0.0059). At 21°C and 26°C, contracture amplitude was unchanged compared to 11°C (for 21°C to 11°C, Holm-corrected p = 0.85; for 26°C to 11°C, Holm-corrected p = 0.21). Thus, while temperature influenced the amplitude of high potassium evoked contractures, the muscles were still capable of contracting at 26°C. In contrast with the semi-intact preparations used for STG-evoked recording, by 20°C, 3 out of 7 preparations had lost contractions, while the remaining 4 had lost the ability to produce contractions by 23°C.

Taken together, this suggests that the contractile machinery of the muscles remains functional at warm temperatures at which other crucial components of the neuromuscular system have failed.

## Discussion

Living systems face a fundamental physical problem: every protein, every ion channel, every membrane, and every biochemical reaction responds to temperature, yet the organism must maintain coordinated function across a wide range of environmental conditions. The components that together produce behavior - from the neurons that generate a motor pattern, to the synapses that relay commands, to the muscles that execute movement - each carry their own individual thermal sensitivity. What is remarkable is that evolution has produced systems capable of sustaining coherent output despite these differences. Rather than making all parts equally robust, or equally temperature sensitive, evolution appears to have built systems in which compensatory changes at one level offset limitations at another, and in which the architecture of the pathway itself distributes the thermal challenge across multiple components working in concert.

Understanding which components are most vulnerable, which provide compensation, and how the system as a whole sustains performance is one of the deep questions in physiology (Robertson and Money, 2012; Marder *et al*., 2015; Marder and Rue, 2021). Thermal resilience is also necessary in the human nervous system, where heat stress, fever, and neurological disease all challenge the coordinated operation of neurons, synapses, and muscles (Kim and Connors, 2012; Wang *et al*., 2014; Thompson, 2015; Peters *et al*., 2016; Christogianni *et al*., 2018; Mavashov *et al*., 2023). Gaining insight, as we do here, into how temperature sensitivity and compensation operate across levels of organization should illuminate temperature resilience in both healthy and neurologically compromised humans.

We addressed this challenge in the neuromuscular transform of *C.borealis*. While previous work has studied the effect of temperature on individual components of motor systems at different levels of organization independently, we looked at them in the interacting system. Our data suggest that, in the transformation from motor pattern generation to movement, the neuromuscular junction is one of the least heat resilient components of this system. The decline in muscle function with warming is likely explained by multiple interacting factors: the hyperpolarization of the muscle fibers (Jacquerie *et al*., 2026), which moves them farther from the threshold for excitation–contraction coupling, the reduction in EJP amplitude and decay time constant, leading to the reduction in synaptic integration, which limits temporal summation and decreases the depolarization produced by each motor neuron spike; and, at extreme temperatures, the failure of the nerve itself, consistent with previous results in related systems (Tang *et al*., 2010; Haddad and Marder, 2018; Schapiro *et al*., 2024; Carrier *et al*., 2026). Together, these effects indicate that temperature does not act on a single isolated process but instead reshapes the entire pathway linking neural activity to contraction.

Muscle fiber hyperpolarization is likely caused by the activation of specific set of potassium currents. Our recent work connects this electrophysiological process with molecular data showing that *C. borealis* expresses two two-pore potassium (K2P) channel genes, *CbKCNK1* (Accession #KU681438) and *CbKCNK2* (Accession #KU681437), homologous to mammalian *KCNK3* and *KCNK9* (Jacquerie *et al*., 2026). This provides a possible molecular explanation for a physiological phenomenon that had been observed previously but had not been linked to specific ion-channel candidates. In this context, the temperature-dependent opening of K2P channels, leading to hyperpolarization and decreases in input resistance, may be advantageous. Because motor nerve activity nearly doubles as temperature increases (Tang *et al*., 2010), the hyperpolarization of the muscle fibers may act as a protective mechanism, preventing excessive depolarization and over-contractability when high-frequency input reaches the muscle. Jointly, reduced input resistance decreases the time constant of EJPs and synaptic integration, which may act to prevent excessive depolarization by reducing summation between spikes (Philip J Stephens, 1985).

This interpretation is supported by the observation that muscle contractility evoked by high-potassium application appears less temperature-sensitive than contractions elicited by nerve stimulation or contractions when the ganglion remains attached to its target muscles. This suggests that the contractile machinery itself may retain substantial function across temperature, whereas synaptic transmission, muscle excitability, or presynaptic release may impose stronger limits on movement. The neuromuscular junction may therefore represent a critical site where temperature-dependent compensation can succeed or fail. More work will be needed to determine how warming affects the presynaptic terminal, transmitter release, synaptic reliability, and the ability of neuromodulators to stabilize neuromuscular output.

In their natural environment, *C.borealis* experience seasonal temperatures that can approach or exceed 20°C. Our experiments, performed in animals acclimated to 11°C, suggest that this mid-range acclimation is not sufficient to preserve robust function across the full thermal range of *C.borealis* experience in the ocean. This implies that long-term acclimation, neuromodulation, or both are likely required to extend the physiological operating range of the system (Blundon, 1989; Chung and Webster, 2004; Zhurov and Brezina, 2005; Hamilton *et al*., 2007; Jury and Watson, 2013; Thuma *et al*., 2013; Lewis and Ayers, 2014). Consistent with this idea, Kedia *et al*. showed that acclimation can shift the operating range of *Homarus americanus* (Kedia *et al*., 2026) and that *C.borealis* network behavior is also influenced by acclimation and ocean experience (Tang *et al*., 2010; Marder and Rue, 2021; Stein *et al*., 2023). A central question for future work is whether acclimation shifts all components of the motor pathway coordinately, or whether some components are disproportionately vulnerable. Does acclimation strengthen the neuromuscular junction, shift circuit activity, alter muscle excitability, or change these processes in parallel? Understanding the relative resilience of each component will be essential for determining how the whole motor system maintains function across environmental temperature variation.

In addition to cellular and molecular processes, behavioral strategies can also help animals remain within a functional thermal range. Migration, habitat selection, or movement into colder microenvironments may allow animals to avoid temperatures that exceed their physiological limits (Kearney, Shine and Porter, 2009). For example, intertidal crabs may also burrow into the sand to reach colder regions, reducing their exposure to extreme temperatures (Watson *et al*., 2018; Hews *et al*., 2021). These behaviors should be considered part of the broader repertoire of temperature compensation, acting together with molecular, cellular, and circuit-level mechanisms to preserve movement and survival (Michaiel and Bernard, 2022).

Many foundational studies on temperature and neuromuscular physiology were performed between 1950 to 1990 (Harri and Florey, 1977; Fischer and Florey, 1981; Stephens and Atwood, 1982, 1983; Philip J. Stephens, 1985). This earlier work established important principles, and, in many cases, our results validate its major conclusions. Then the field moved on. Between roughly 1990 and 2020, studies relevant to the effects of temperature and other environmental conditions on neuromuscular junction and muscle excitability in non-model organisms have largely disappeared from the literature. Today, we can return to some of these questions in the context of the challenges posed by altered environmental factors. What was missing then - and what we now have - is the ability to move from the phenomenological to the mechanistic. We can sequence genomes, identify ion channel families, relate molecular expression to cellular physiology, and connect single-channel biophysics to whole-system behavior.

The urgency of this work extends beyond a single crustacean species or a neuromuscular preparation. As environmental temperatures rise at unprecedented rates due to anthropogenic changes, animals are being forced to operate closer to the limits of their physiological capacity. Yet we still know too little about the cellular and biophysical mechanisms that determine whether a nervous system, a synapse, or a muscle will remain functional under thermal stress. If neuroscience is to contribute to our understanding of climate resilience, it must re-engage with fundamental physiology in ecologically relevant organisms, such as marine crustaceans. Ion channels, synapses, muscles, and behavior are the linked mechanisms through which animals either adapt to environmental change or fail to do so (Michaiel and Bernard, 2022). Revisiting these questions with modern tools is therefore essential for building a predictive understanding of how nervous systems and the behaviors they generate will respond to a rapidly changing world.

## Materials and methods

### Animal

Adult male Jonah Crabs (*Cancer borealis*) were purchased from Commercial Lobster Company (Boston, MA) and held in filtered, circulating, aerated, artificial seawater (Instant Ocean Sea Salt Mix) at 11°C in a 12h light/ 12h dark cycle. Crabs were acquired in April 2025, then between September 2025 and March 2026. Animals were typically held in the laboratory for one week before use.

### Experimental preparation

Stomach musculature and the stomatogastric nervous system (STN) were dissected following the description and nomenclature of (Maynard and Dando, 1974). Nerve dissection followed the procedures described by (Gutierrez and Grashow, 2009). The fat layer and connective tissue overlaying the muscles were carefully removed to improve access for recording from muscle fibers while keeping nerve terminals attached (Blitz *et al*., 2017). Reduced preparations of the muscles and nerves were pinned into a silicone elastomer-lined (Sylgard 184: Dow Corning) dish in physiological saline. The *C. borealis* saline solution (in mM): 440 NaCl, 11 KCl, 26 MgCl_2_, 13 CaCl_2_, 11 Trizma base, 5.4 maleic acid, pH 7.8 at 11°C. Solutions were calibrated using a Mettler Tolledo pH meter. High-potassium saline (10xK^+^) was adjusted to 110 mM KCl, and all other components remained the same. Saline was continuously superfused at 8-11 mL/min via a peristaltic pump (model Ismatec Ecoline). For the high-potassium experiment, the flow rate was at 25 mL/min. Temperature was adjusted from 6°C to 26°C using a Peltier system (Warner Instruments SC-20) with a temperature controller (CL-100, Warner Instruments) and a thermocouple probe in the bath near the recording site. A tolerance interval of ± 0.3°C of the target value (e.g., 10.7 - 11.3°C for 11°C) was considered acceptable.

### Electrophysiology

Electrophysiology experiments were performed as previously described (Jorge-Rivera *et al*., 1998; Daur *et al*., 2012; Blitz *et al*., 2017; Daur, Nadim and Bucher, 2021; Jacquerie *et al*., 2026). For multi-level recording, the STN was intact, including the paired commissural ganglia, the esophageal ganglion, and the stomatogastric ganglion (STG). For simultaneous recording of STG, EJP and contraction, all muscles were discarded except the LP-innervated muscle (cpv4, cpv6, and p1) (Figure 1D). For simultaneous recording of STG and EJP, PY-innervated muscle (p2) was also kept. The lvn was recorded using a suction electrode. Signals were amplified using a 16-channel AC amplifier (Model 3500; AM Systems).

Intracellular recordings of EJPs in muscle fibers were obtained using sharp borosilicate glass microelectrodes (Sutter Instrument; ID 0.86 mm, OD 1.5 mm), pulled with a micropipette puller (Sutter Instrument, P-97) and backfilled with 400 mM K_2_S0_4_, 20 mM KCl, and 33 mM NaCl. Electrode resistances were between 5 MΩ and 20 MΩ. Electrodes were mounted on Leica Leitz mechanical micromanipulators with HS-2A-x1LU Axoclamp 2B and HS-9A-x1U Axoclamp 900A headstages (Molecular Devices). Signals were amplified using Axoclamp 2B or 900A amplifiers (Molecular Devices), digitized at 10 kHz using a Digidata 1440, and recorded in Clampe× 10.7. The electrodes had an offset between 0 and 3mV for a 10°C increase from 11°C to 21°C.

Force generated by the muscle was recorded using a force-displacement transducer (FT03, Grass Instruments). One muscle insertion was pinned down while the other side was sutured to the force-displacement transducer held by a clamp at 45°. The muscles were stretched to their resting length. The force-displacement transducer was calibrated by applying a known weight corresponding to a force of 1 cN, allowing conversion of the recorded voltage signal to force units.

The nerve was stimulated using a suction electrode attached to the cut nerve end, connected to an AM Systems isolated pulse stimulator (Model 2100). Pulse trains were delivered mimicking the realistic activity of a LP neuron or paired-pulse stimuli. Individual pulse durations ranged from 0.2 ms to 0.9 ms. Pulse amplitudes were calibrated to reliably trigger contraction or a single EJP at 11°C while keeping a 10 - 20% safety margin. Muscle fibers having a resting membrane voltage beyond −45 mV were excluded from recordings (Daur *et al*., 2012).

The current versus voltage plot was recorded in two-electrode current clamp mode. Two electrodes were inserted in the same fiber along the longitudinal axis. The distance between the two electrodes was less than the diameter of the fiber. Current-voltage (I-V) curves were obtained by stepping the current each 10 nA for 2 s such that the voltage range explored was between −100 mV and −50 mV. Saline levels were minimized to reduce capacitive coupling. The headstage for recording currents was a Molecular Devices HS-2A-x10MU Axoclamp 2B, connected via an adapter to the Axoclamp 900A. Because the muscle fibers are large and electrically coupled, and produce substantial currents, current steps were accepted only if they reached the intended values and the recorded currents remained stable. The electrode used for current injection was filled with 3 M KCl.

### Quantification

We developed custom MATLAB and Python scripts to visualize and analyze recordings obtained using Clampex software. Figures were made in Adobe Illustrator.

<u>LP spikes</u> were detected and sorted from extracellular lvn recordings using Crabsort (Gorur-Shandilya *et al*., 2022). LP bursts, LP spikes per burst, LP duty cycle, and LP intraburst spike frequency were used to quantify pyloric activity. Measurements were assigned to temperature bins centered at 11, 14, 17, 20, and 23°C, using a half-width of 1.3°C around each bin center. For each preparation and temperature bin, we calculated the median value of each LP feature. Group values are reported as mean ± standard error of the mean (SEM) across preparations.

To examine the temperature dependence of the <u>EJPs evoked by STG</u> <u>activity</u> (Figures 1D-E), for each muscle, EJP amplitude was defined as the difference between peak membrane depolarization and baseline membrane potential. The amplitudes were assigned to temperature bins centered at 11, 14, 17, 20, and 23°C, using a half-width of 1.3°C around each bin center. For each animal, the median EJP amplitude was calculated within each temperature bin. We averaged the per-animal medians across animals for each muscle and bin. Error bars represent the SEM across animals.

For Figure 2E, <u>force recordings</u> were converted to mN using the calibration factor determined for each experiment. For each pyloric cycle, the contraction amplitude was defined as the difference between the local peak force and the preceding local minimum force. For each preparation and temperature bin, contraction amplitude was summarized as the median contraction amplitude across cycles. Group values were then calculated across preparations.

For Figure 2F, a <u>successful contraction</u> was defined as a preparation in which pyloric nerve activity remained rhythmic, the force signal showed visible, cycle-locked contractions, and the contraction amplitude was at least 15% of that measured at 11°C in the same preparation. Preparations in which rhythmic pyloric nerve activity persisted but force amplitude fell below threshold were classified as having lost effective contractile function.

For <u>nerve-evoked contractions</u> in Figure 3, the maximal force was measured as the peak force reached during the stimulation train relative to the pre-stimulation baseline. To avoid scoring noise or flat traces as contractions, peaks were accepted only when their amplitude exceeded an adaptive threshold based on the baseline noise of each recording. The minimum accepted peak amplitude was defined as the larger of 2 mV or 2.5 times the baseline noise estimate, with an upper cap of 50 V for the adaptive threshold. Peaks below this threshold were treated as absent.

To quantify the relative <u>phasic and tonic components of the contraction</u>, we analyzed the response to the final burst of the stimulation train, following the logic of previous analyses of crustacean stomach muscle contractions (Morris and Hooper, 1998; Morris, Thuma and Hooper, 2000). The total contraction amplitude was defined as the force reached during the final burst relative to the pre-stimulation baseline. The phasic component was defined as the oscillatory force component produced by the final burst, measured relative to the force level immediately preceding that burst. The tonic component was therefore the sustained force remaining at the start of the final burst. Percent phasic amplitude was calculated as: 100 × phasic amplitude / total contraction amplitude. When the force trace had returned to baseline before the final burst, the tonic component was interpreted as 0, and the percent phasic amplitude was set to 100%. For each preparation, temperature, and stimulation pattern, values were first summarized within preparation and then averaged across preparations. Group values are reported as mean ± SEM.

For each nerve-evoked recording, <u>the relaxation time constant</u> was extracted from the contraction produced by the final burst. The baseline was defined as the mean force during the final 1 s of the recording, and the peak amplitude of the last contraction was measured relative to this baseline. The half-relaxation point was then identified as the time after the peak when the force decayed to halfway between the peak and baseline. A mono-exponential decay was fit from this half-relaxation point to 1 s later, using the form *_F_*(*_t_*) = *_Ae_*^!“/$^, and the fitted *_τ_* value was taken as the relaxation time constant. Traces with contraction amplitudes below 0.3 mN were excluded from the quantification.

For <u>paired-pulse experiments</u> in Figure 4, EJPs were analyzed from intracellular recordings of cpv4 during lvn stimulation at different paired-pulse frequencies. For each burst, the resting membrane potential was defined as the mean membrane voltage during the 100 ms window preceding stimulation onset. Stimulation artifacts were removed from the voltage traces before peak detection. EJP peaks were then detected after the stimulation artifact. <u>EJP amplitude</u> was measured relative to the pre-stimulation resting membrane potential. For each paired-pulse response, the first EJP amplitude was defined as the amplitude of the first peak relative to rest. The second EJP amplitude was defined as the peak membrane depolarization reached after the second stimulus relative to rest.

The <u>decay time constant</u> of the first EJP was extracted from the falling phase of the first EJP. The decay segment began at the first EJP peak and extended until the local minimum before the second EJP, or to the end of the analysis window when no subsequent peak was present. The decay was fit with an exponential function, and the fitted time constant was used as the EJP decay time constant. For Figure 4G, the time constant of the first EJP obtained of the 1 Hz stimulus was used for statistical comparison across temperatures.

<u>Summation</u> was quantified as the residual depolarization remaining from the first EJP at the time immediately before the second EJP. For low and intermediate stimulation frequencies, this value was measured as the minimum membrane potential between the first and second EJP peaks, relative to the resting membrane potential. For high stimulation frequencies, where the second EJP occurred before the first EJP had fully decayed, the residual depolarization from the first EJP was estimated from the fitted decay. This residual component was subtracted when necessary to estimate the net amplitude of the second EJP. Absolute summation was reported in mV. Percent summation was calculated as: 100 × summation / EJP_2_ amplitude, where EJP_2_ amplitude was the second EJP peak measured relative to the resting membrane potential. For each animal, temperature, and stimulation frequency, different fiber recordings were averaged before group statistics were calculated on animals.

<u>Input resistance (R_in_)</u> was estimated from current-voltage relationships obtained during two-electrode current-clamp recordings. For each sweep, voltage and current were measured by averaging the signals over a 10 ms window centered at 2.5 s of the step to be in steady-state. One voltage-current point was extracted per sweep. R_in_ was then calculated for each file by fitting a linear regression of membrane voltage as a function of injected current using all finite data points with current values ≤ 10 nA. Input resistance was computed from the slope of the fit. The goodness of fit was assessed using the coefficient of determination (R^2^)

For <u>high-potassium experiments</u>, contractures were evoked by applying 10xK^+^ saline for 40 s at 11, 16, 21, and 26°C. Muscles were allowed to recover for 30 min between applications before the temperature was increased for the next trial. Contracture amplitude was quantified as the maximal force reached during the 10xK^+^ application relative to the baseline force immediately preceding the solution switch. Baseline force was calculated from a stable pre-application window before the onset of the contracture. For each preparation and temperature, the peak contracture amplitude was extracted and used for statistical analysis. Group values are reported as mean ± SEM across preparations.

### Statistical analysis

For statistical analyses, data are reported as either the median or the mean ± SEM, as indicated in the figure legends. Significant differences are indicated as follows: *p < 0.05, **p < 0.01, ***p < 0.001.

Temperature effects on LP motor features in Figure 2A-D were tested separately for each feature using linear mixed-effects models with temperature as a categorical fixed effect and preparation identity as a random intercept: feature value ∼ temperature + (1 | preparation). When the omnibus temperature effect was significant, post hoc test comparisons were made between each temperature and 11°C using model contrasts, with Holm correction for multiple comparisons within each feature. Details about the number of animals and the p-values are reported in Table S1 and in Table S2.

For Figure 2E, maximal EJP amplitude was analyzed using a linear mixed-effects model with temperature, muscle identity, and their interaction as categorical fixed effects, and preparation identity as a random intercept: Max EJP amplitude ∼ temperature × muscle + (1 | preparation). Follow-up within-muscle comparisons against 11°C were performed using paired Wilcoxon signed-rank tests after averaging multiple recordings from the same preparation, with Holm correction within each muscle. Number of animals are reported in Table S3.

For Figure 3B<u>, nerve-evoked contraction amplitude</u> was analyzed using a linear mixed-effects model with temperature, stimulation pattern, and their interaction as categorical fixed effects, and preparation identity as a random intercept: amplitude ∼ temperature × stimulation pattern + (1 | preparation). Degrees of freedom were estimated using Satterthwaite’s approximation. Contraction amplitude was analyzed using a linear mixed-effects model with temperature, stimulation pattern, and their interaction as categorical fixed effects, and preparation identity as a random intercept: amplitude ∼ temperature × stimulation pattern + (1 | preparation). Degrees of freedom were estimated using Satterthwaite’s approximation. After fitting the model, we performed pre-specified comparisons to test whether the fastest LP-like stimulation pattern increased force at 21°C relative to slower patterns, and whether this increase restored force to the levels observed at cooler temperatures. Holm correction was applied within each set of comparisons. One-sided tests were used only for incomplete-rescue comparisons, for which we had the a priori prediction that cooler reference conditions would produce greater force than the LP21 pattern at 21°C. Group summaries are shown as the mean ± SEM across preparations.

Table S4 reports the number of animals for each temperature and stimulus pattern used for the phasic amplitude analysis. Table S5 reports the number of animals for each temperature and stimulus pattern.

For Figure 4B, EJP_2_ amplitude was analyzed using a linear mixed-effects model with stimulation frequency, temperature, and their interaction as categorical fixed effects, and recording identity as a random intercept: EJP_2_ amplitude ∼ temperature × frequency + (1 | recording). Stimulation frequency had six levels: 1, 5, 10, 20, 30, and 40 Hz, with 1 Hz as the reference. Temperature had three levels: 11, 16, and 21°C, with 11°C as the reference. We performed the statistical comparison on individual fibers. We had 7 fibers from 5 animals. EJP_2_ amplitude was defined as the peak membrane depolarization reached after the second stimulus, measured relative to the resting membrane potential immediately preceding stimulation onset.

The significance of each fixed effect was assessed from the model ANOVA table. Three families of post hoc pairwise comparisons were then computed from the model-estimated means and their covariance matrix, with Holm correction applied within each family.

First, to test whether higher stimulation frequencies produced larger EJP₂ amplitudes than the lowest frequency, EJP_2_ at each frequency was compared against EJP_2_ at 1 Hz within each temperature (one-sided, H_1_: amplitude at higher frequency > amplitude at 1 Hz).

Second, to test whether warming reduced EJP_2_₂ amplitude, each frequency condition at 16°C and 21°C was compared against the same frequency at 11°C (one-sided, H_1_: amplitude at warm temperature < amplitude at 11°C).

Third, to test whether high-frequency paired-pulse stimulation at warm temperature could compensate for the temperature-dependent reduction in EJP_2_ amplitude, EJP_2_ at 40 Hz and 21°C was compared against EJP₂ at 10 Hz at 11°C and against EJP_2_ at 20 Hz at 11°C (one-sided, H_1_: 40 Hz at 21°C > reference condition at 11°C; Holm correction across the two comparisons). This tests whether the motor circuit’s temperature-dependent increase in firing frequency can offset the synaptic depression produced by warming. Detailed values are reported in Table S6.

For Figure 4C, the effect of temperature on resting membrane potential was tested using a repeated-measures Friedman test across 11, 16, and 21°C. Multiple fibers from the same animal were averaged so the unit of analysis was the animal.

For Figure 4E, the effect of temperature on the input resistance was tested using a repeated-measure Friedman test across 11, 14, 17, and 20°C.

For Figure 4F, the effect of temperature on the decay time constant of the first EJP was tested using a repeated-measures Friedman test across 11, 16, and 21°C. Multiple fibers from the same animal were averaged so the unit of analysis was the animal.

For Figure 5B, the effect of temperature on high-potassium contracture amplitude was analyzed using a repeated-measures Friedman test across 11, 16, 21, and 26°C, with each animal as the unit of analysis. Only preparations with complete measurements at all four temperatures were included. When the overall Friedman test was significant, three pairwise post-hoc comparisons were performed between each warmer temperature (16, 21, and 26°C) and the 11°C reference using two-sided Wilcoxon signed-rank tests, with Holm correction across the three comparisons.

## Acknowledgements

We thank Sonal Kedia, Kyra Schapiro, and Natasha Baas-Thomas for their feedback on this project; Gwen Harris and Alice Lander for the management of the animals and laboratory facility; Scott Hooper and Christoph Guschlbauer for their insightful feedback; Francisco Mello Jr. for his help with equipment.

## Funding

This work was supported by the United States National Institute of Health R35 NS 142987 (E.M.; J.M.D; J.Z.), R01 MH 046742 (J.Z.), Computational Neuroscience Training Grant 5T90DA059112-03 (J.M.D); Belgian American Educational Foundation - BAEF, Wallonie- Belgium International - WBI Postdoctoral Fellowship (K.J.).

## Notes

### Competing Interest Statement

The authors have declared no competing interest.

