## Supplementary Information for "Compensation of the effects of temperature on a motor system in the crab, *Cancer borealis*"

### Table S1: Number of animals for Figure 1 A-D

| Feature\ n | 11°C | 14°C | 17°C | 20°C | 23°C |
|---|---|---|---|---|---|
| **Spikes per burst** | 13 | 14 | 14 | 14 | 9 |
| **Spike frequency** | 13 | 14 | 14 | 14 | 9 |
| **Pyloric frequency** | 13 | 14 | 14 | 14 | 9 |
| **Duty cyle** | 13 | 14 | 14 | 14 | 9 |

### Table S2: Statical table for Figure 1 A-D

| Feature | Comparison | n 11°C | n test | 11°C mean ± SEM | Test mean ± SEM | Signf. | Direction |
|---|---|---|---|---|---|---|---|
| **Spikes per burst** | 14°C vs 11°C | 13 | 14 | 6.27 ± 0.74 | 6.36 ± 0.72 | n.s. | increase vs 11°C |
| **Spikes per burst** | 17°C vs 11°C | 13 | 14 | 6.27 ± 0.74 | 6.00 ± 0.68 | n.s. | decrease vs 11°C |
| **Spikes per burst** | 20°C vs 11°C | 13 | 14 | 6.27 ± 0.74 | 4.64 ± 0.68 | *** | decrease vs 11°C |
| **Spikes per burst** | 23°C vs 11°C | 13 | 9 | 6.27 ± 0.74 | 3.89 ± 0.75 | *** | decrease vs 11°C |
| **Spike frequency (Hz)** | 14°C vs 11°C | 13 | 14 | 24.08 ± 2.22 | 29.69 ± 2.85 | * | increase vs 11°C |
| **Spike frequency (Hz)** | 17°C vs 11°C | 13 | 14 | 24.08 ± 2.22 | 33.88 ± 3.03 | *** | increase vs 11°C |
| **Spike frequency (Hz)** | 20°C vs 11°C | 13 | 14 | 24.08 ± 2.22 | 38.00 ± 3.87 | *** | increase vs 11°C |
| **Spike frequency (Hz)** | 23°C vs 11°C | 13 | 9 | 24.08 ± 2.22 | 44.12 ± 3.88 | *** | increase vs 11°C |
| **Pyloric frequency (Hz)** | 14°C vs 11°C | 13 | 14 | 0.97 ± 0.06 | 1.21 ± 0.09 | * | increase vs 11°C |
| **Pyloric frequency (Hz)** | 17°C vs 11°C | 13 | 14 | 0.97 ± 0.06 | 1.44 ± 0.11 | *** | increase vs 11°C |
| **Pyloric frequency (Hz)** | 20°C vs 11°C | 13 | 14 | 0.97 ± 0.06 | 1.77 ± 0.11 | *** | increase vs 11°C |
| **Pyloric frequency (Hz)** | 23°C vs 11°C | 13 | 9 | 0.97 ± 0.06 | 2.26 ± 0.19 | *** | increase vs 11°C |
| **Duty cycle (%)** | 14°C vs 11°C | 13 | 14 | 20.72 ± 1.84 | 20.92 ± 1.93 | n.s. | decrease vs 11°C |
| **Duty cycle (%)** | 17°C vs 11°C | 13 | 14 | 20.72 ± 1.84 | 20.34 ± 1.99 | n.s. | decrease vs 11°C |
| **Duty cycle (%)** | 20°C vs 11°C | 13 | 14 | 20.72 ± 1.84 | 15.19 ± 2.55 | *** | decrease vs 11°C |
| **Duty cycle (%)** | 23°C vs 11°C | 13 | 9 | 20.72 ± 1.84 | 13.28 ± 3.12 | *** | decrease vs 11°C |

### Table S3: Statistical table for Figure 2E

| n | 11°C | 14°C | 17°C | 20°C | 23°C |
|---|---|---|---|---|---|
| **p1** | 5 | 7 | 7 | 7 | 4 |
| **p2** | 6 | 9 | 9 | 9 | 7 |
| **cpv4** | 11 | 11 | 12 | 12 | 6 |

Table S4: Statical table for Figure 3

| Question | Comparison | Estimate (mN) | p-value | Adjusted p-value | significance |
|---|---|---|---|---|---|
| **Does temperature affect force?** | Temperature | — | $9.82 \times 10^{-5}$ | — | *** |
| **Does stimulation pattern affect force?** | Stimulation | — | 0.038 | — | * |
| **Interaction** | Temperature x simulation | — | 0.315 | — | n.s. |
| **Rescue at 21°C** | LP21 − LP05 | 2.97 | 0.0057 | 0.019 | * |
| **Rescue at 21°C** | LP21 − LP11 | 2.44 | 0.0047 | 0.019 | * |
| **Rescue at 16°C** | LP16 − LP05 | 2.88 | 0.00021 | 0.001 | ** |
| **Incomplete rescue** | LP21: 21°C − 11°C | −5.00 | 0.007 | 0.019 | * |
| **Temperature decline at low frequency** | LP05: 21°C 11°C | −4.08 | 0.0076 | 0.019 | * |
| **Is LP 11 at 11°C greater than the LP 21 at 21°C** | 11°C LP 11 – 21°C LP 21 | 3.56 | 0.0020 | 0.006 | ** |

Table S4: Number of animals for Figure 3C

| n | 6°C | 11°C | 16°C | 21°C |
|---|---|---|---|---|
| **LP 6** | 7 | 7 | 7 | 4 |
| **LP 11** | 7 | 7 | 7 | 6 |
| **LP 16** | 7 | 7 | 7 | 6 |
| **LP 21** | 7 | 7 | 7 | 7 |

Table S5: Number of animals for Figure 3D

| n | 6°C | 11°C | 16°C | 21°C |
|---|---|---|---|---|
| **LP 6** | 7 | 7 | 4 | 0 |
| **LP 11** | 7 | 7 | 6 | 3 |
| **LP 16** | 7 | 7 | 7 | 4 |
| **LP 21** | 7 | 7 | 7 | 5 |

Table S6: Statistical table for Figure 4B

| Question | Test / comparison | Estimate | p value | Holm-corrected p | Significance |
|---|---|---|---|---|---|
| **Overall effect of stimulation condition** | LME fixed effect | F = 21.4 | $7.59 \times 10^{-15}$ | — | **** |
| **Overall effect of temperature** | LME fixed effect | F = 4.4 | 0.015 | — | * |
| **Stimulation × temperature interaction** | LME fixed effect | F = 0.67 | 0.75 | — | n.s. |
| **$EJP_2$ at 1 Hz at 21°C vs 11°C** | 21°C − 11°C | −3.80 mV | 0.00341 | 0.024 | * |
| **$EJP_2$ at 5 Hz at 21°C vs 11°C** | 21°C − 11°C | −4.33 mV | $1.08 \times 10^{-3}$ | $8.64 \times 10^{-3}$ | ** |
| **$EJP_2$ at 10 Hz at 21°C vs 11°C** | 21°C − 11°C | −5.28 mV | $1.08 \times 10^{-4}$ | $1.04 \times 10^{-3}$ | ** |
| **$EJP_2$ at 20 Hz at 21°C vs 11°C** | 21°C − 11°C | −5.30 mV | $1.04 \times 10^{-4}$ | $1.04 \times 10^{-3}$ | ** |
| **$EJP_2$ at 30 Hz at 21°C vs 11°C** | 21°C − 11°C | −7.17 mV | $4.75 \times 10^{-7}$ | $5.70 \times 10^{-5}$ | *** |
| **$EJP_2$ at 40 Hz at 21°C vs 11°C** | 21°C − 11°C | −6.59 mV | $2.82 \times 10^{-6}$ | $3.10 \times 10^{-5}$ | *** |
| **Comparison test** | 21°C $EJP_2$ 40 Hz > 11°C $EJP_2$ 10 Hz | +0.65 mV | 0.640 | 0.641 | n.s. |
| **Comparison test** | 21°C $EJP_2$ 40 Hz > 11°C $EJP_2$ 20 Hz | −2.35 mV | 0.092 | 0.184 | n.s. |

Table S7: Statistical table for Figure 5

| Question | Test | Comparison | Mean ≠ (mN) | Median ≠ (mN) | Raw p | Holm-cor. p | Significance |
|---|---|---|---|---|---|---|---|
| **Does cpv46p1 force change with temperature?** | Friedman test | 11, 16, 21, 26°C | — | — | 0.00023 | — | *** |
| **Does force differ from 11°C?** | Wilcoxon signed-rank, two-sided | 16°C vs 11°C | 26.6 | 28.5 | 0.0020 | 0.0058 | ** |
| **Does force differ from 11°C?** | Wilcoxon signed-rank, two-sided | 21°C vs 11°C | 2.11 | 8.65 | 0.85 | 0.85 | n.s. |
| **Does force differ from 11°C?** | Wilcoxon signed-rank, two-sided | 26°C vs 11°C | −18.3 | −25.7 | 0.105 | 0.21 | n.s. |
